# Multicellular Perturb-seq Reveals Lineage-Specific Transcriptional Control During Human Cardiac Organoid Development

**DOI:** 10.64898/2026.11.05.763441

**Authors:** Payton Klosa, Kuancheng Cai, Coneria Nansubuga, Yifan Sun, Xueqiu Lin, Chun Liu

## Abstract

Heart development requires the coordinated differentiation of multiple cell lineages, including cardiomyocytes (CMs) and endothelial cells (ECs). Transcription factors (TFs) orchestrate lineage specification, morphogenesis, and maturation, yet how cardiac and vascular TFs shape multicellular organization and their interactions remain poorly understood. Many TFs exhibit pleiotropic and stage-dependent effects across cardiac lineages, particularly in cardiac progenitor cells, suggesting the presence of shared regulatory networks that synchronize cardiac and vascular programs during development and in response to stress. Human induced pluripotent stem cell (hiPSC) differentiated cardiac organoid provides a physiologically relevant 3D model to study these multicellular interactions, enabling simultaneous differentiation of ECs and CMs within a human genetic context. When combined with CRISPR interference (CRISPRi)–based Perturb-seq, this system allows systematic, high-throughput dissection of TF-driven regulatory programs at single-cell resolution. In this study, we investigated how cardiac and vascular TFs coordinate lineage commitment and the cross-lineage signaling in human cardiovascular differentiation.

## Introduction

Key cardiac TFs, including GATA4, HAND1, HAND2, MEF2C, NKX2-5, and TBX5, and vascular regulators, including ETV2, KLF2, NOTCH1, and SOX17, play critical roles in cardiovascular development (Srivastava 2006; Grunert et al. 2024). However, individual TFs often exhibit stage-dependent and lineage-specific functions, and how these regulatory programs converge or diverge during multicellular cardiovascular differentiation remains poorly understood. Human pluripotent stem cell-derived cardiac organoids provide a promising model to investigate these processes by enabling the simultaneous differentiation and self-organization of cardiac and vascular cell populations within a three-dimensional tissue environment (Voges et al. 2023). Nevertheless, systematic functional interrogation of TF networks across developmental stages and multiple cardiovascular lineages remains limited. Perturb-seq, originally developed by Dixit et al. and Adamson et al. in 2016, integrates pooled CRISPR-mediated genetic perturbations with single-cell RNA sequencing to systematically characterize gene regulatory functions at single-cell resolution (Adamson et al. 2016; Dixit et al. 2016). Recent applications in human brain organoids have demonstrated its ability to identify TF-dependent cell fate decisions, developmental defects, and lineage-specific regulatory networks within complex multicellular systems (Fleck et al. 2023; Li et al. 2023). Furthermore, Perturb-seq has been applied to differentiating human pluripotent stem cells to dissect transcriptional regulatory networks underlying cardiomyocyte and vascular differentiation separately (Nikolova et al. 2025; Sivakumar et al. 2025). However, the stage-dependent requirements of cardiac and vascular TFs during coordinated multicellular cardiovascular differentiation remain incompletely understood. Here, we establish a time-resolved CRISPR interference (CRISPRi) Perturb-seq platform in human cardiac organoids (CO) to systematically investigate ten cardiovascular TFs across four developmental stages. Our analysis identifies distinct stage- and lineage-specific TF requirements, reveals convergent roles of MEF2C, HAND2, and NKX2-5 in cardiac progenitor-to-cardiomyocyte differentiation, and uncovers divergent regulation of mesenchymal transcriptional programs. Together, this study provides a framework for dissecting transcriptional regulation of human cardiovascular development within a multicellular context using organoid.

## Results

### A time-resolved single-cell Perturb-seq analysis of endothelial cell and cardiomyocyte differentiation using cardiac organoid

We differentiated a CRISPRi (dCas9-KRAB) knockin human iPSC line through a optimized cardiac organoid (CO) protocol in which mesoderm induction (D0) is followed by cardiac patterning (D3–D4) as previous published cardiod protocol and gastruloid protocol (Schmidt et al. 2023; Abilez et al. 2025). Day 4 VEGF promotes vascular patterning, then following CO specification (D6) and maintenance (D15), with endpoint analysis at D22 (**Fig. 1A**). COs compacted progressively over the first week and, by D22, contained an interactive CD31^+^ endothelial network wrapped around and through a TNNT2^+^ myocardial core (**Fig. 1A**), confirming that the cardiac and vascular lineages co-differentiate within a single organoid rather than segregating into separate aggregates.

**Figure 1.**
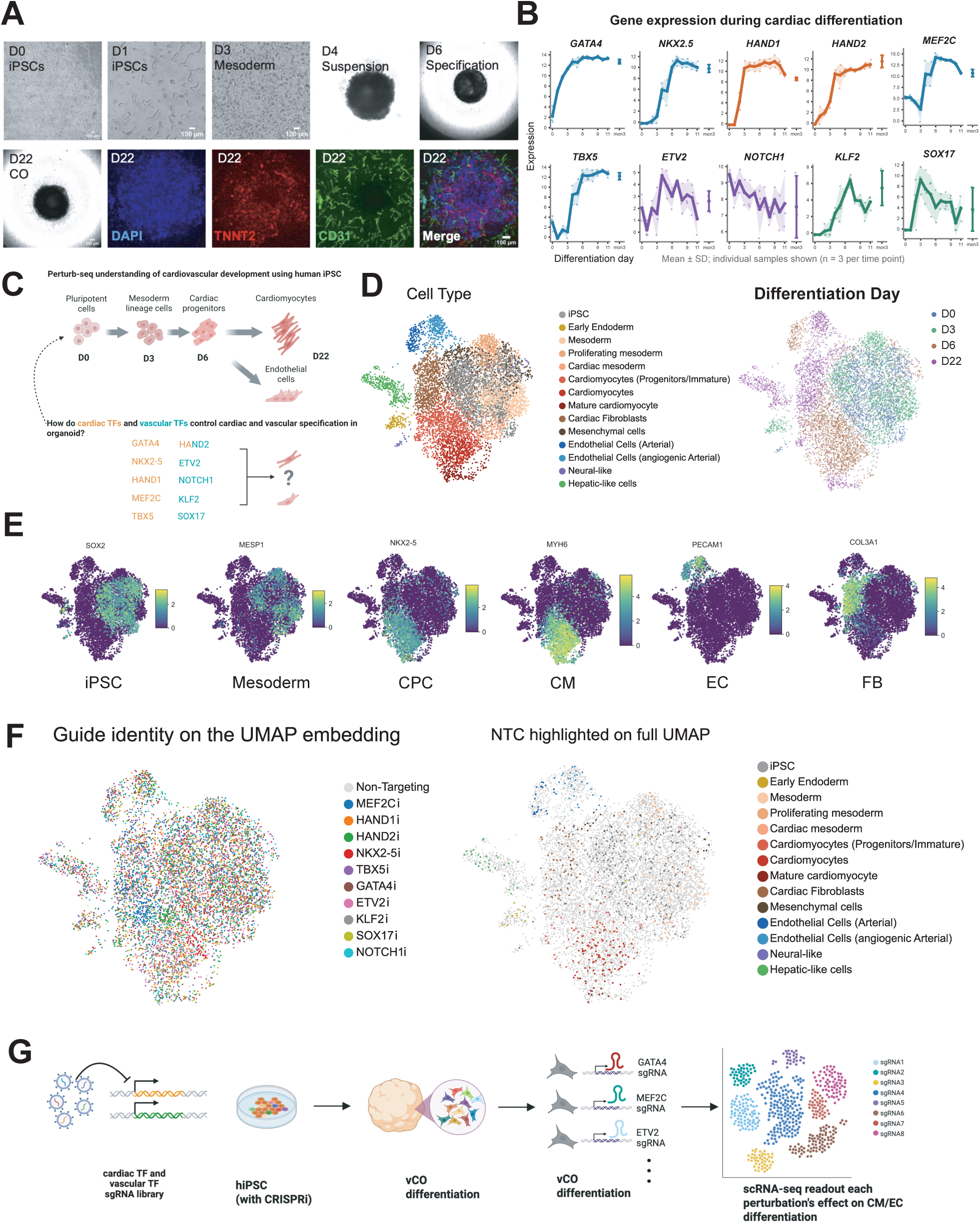
Time-resolved multicellular Perturb-seq during human cardiac organoid differentiation. (A) Cardiac organoid differentiation with representative organoid images. (B) Temporal expression of selected cardiac and vascular transcription factors (TFs) during differentiation. (C) Experimental design and single-cell profiling at D0, D3, D6, and D22. (D) UMAP visualization of 14 identified cell states across differentiation stages. (E) Expression of canonical lineage-specific markers. (F–G) Distribution of sgRNA-assigned cells across developmental stages and cell populations.

We transduced the line with a pooled sgRNA library targeting six important cardiac TFs (GATA4, HAND1, HAND2, MEF2C, NKX2-5, TBX5) and four important vascular TFs (ETV2, KLF2, NOTCH1, SOX17), three sgRNAs per gene, together with three non-targeting controls (NTC) in the CRISPRi iPSC line at 0.2-0.3 MOI. These important TFs were significantly activated around D6 cardiac progenitor stage, validated using previous human iPSC cardiac differentiation time-course bulk RNA-seq data (**Fig. 1B and 1C**). To avoid potential selection effects of antibiotic selection during iPSC expansion after sgRNA library expansion, the transduced iPSCs were not enriched using puromycin selection in sgRNA backbone. Perturbation and single-cell transcriptomes were collected at D0, D3, D6 and D22 according to iPSC stage, mesoderm stage, cardiac progenitor stage, and differentiated CO stage (**Fig. 1C**). After filtering and integration, the atlas resolved fourteen states: iPSC, mesoderm, proliferating mesoderm, cardiac mesoderm, early endoderm, cardiomyocyte progenitor/immature, cardiomyocyte, mature cardiomyocyte, arterial endothelium, angiogenic arterial endothelium, mesenchymal cells, cardiac fibroblasts, hepatic-like cells and neural-like cells (**Fig. 1D**). Collection day was the dominant axis of variation on the embedding (**Fig. 1D**). Canonical markers separated the states as expected: SOX2 and NANOG in iPSC, MESP1 in mesoderm, NKX2-5 and PDGFRA in cardiac progenitor cell (CPC), MYH6, MYH7 and ACTN2 in cardiomyocytes (CM), CDH5 and PECAM1 in endothelium (EC), SERPINA1 in hepatic-like cells and COL3A1 in the fibroblasts (FB) (**Fig. 1E**).

We therefore assigned each cell to its dominant guide, recovering 8,068 guide-assigned cells across the four timepoints: MEF2C 992, HAND2 924, TBX5 837, ETV2 836, GATA4 763, SOX17 726, NTC 726, HAND1 675, NOTCH1 592, NKX2-5 534 (**Fig. 1F and 2A**). Per-guide, per-day coverage ranged from 85 cells (KLF2, D22) to 332 cells (MEF2C, D22), with a median near 190 (**Fig. 1F** and supplementary table 1). TF guides and NTC guides were distributed across the whole embedding for every target and almost captured at every cell type during differentiation (**Fig. 1F and 1G**). Restricting to NTC cells gave the unperturbed reference trajectory for TF perturbation effects analysis.

**Figure 2.**
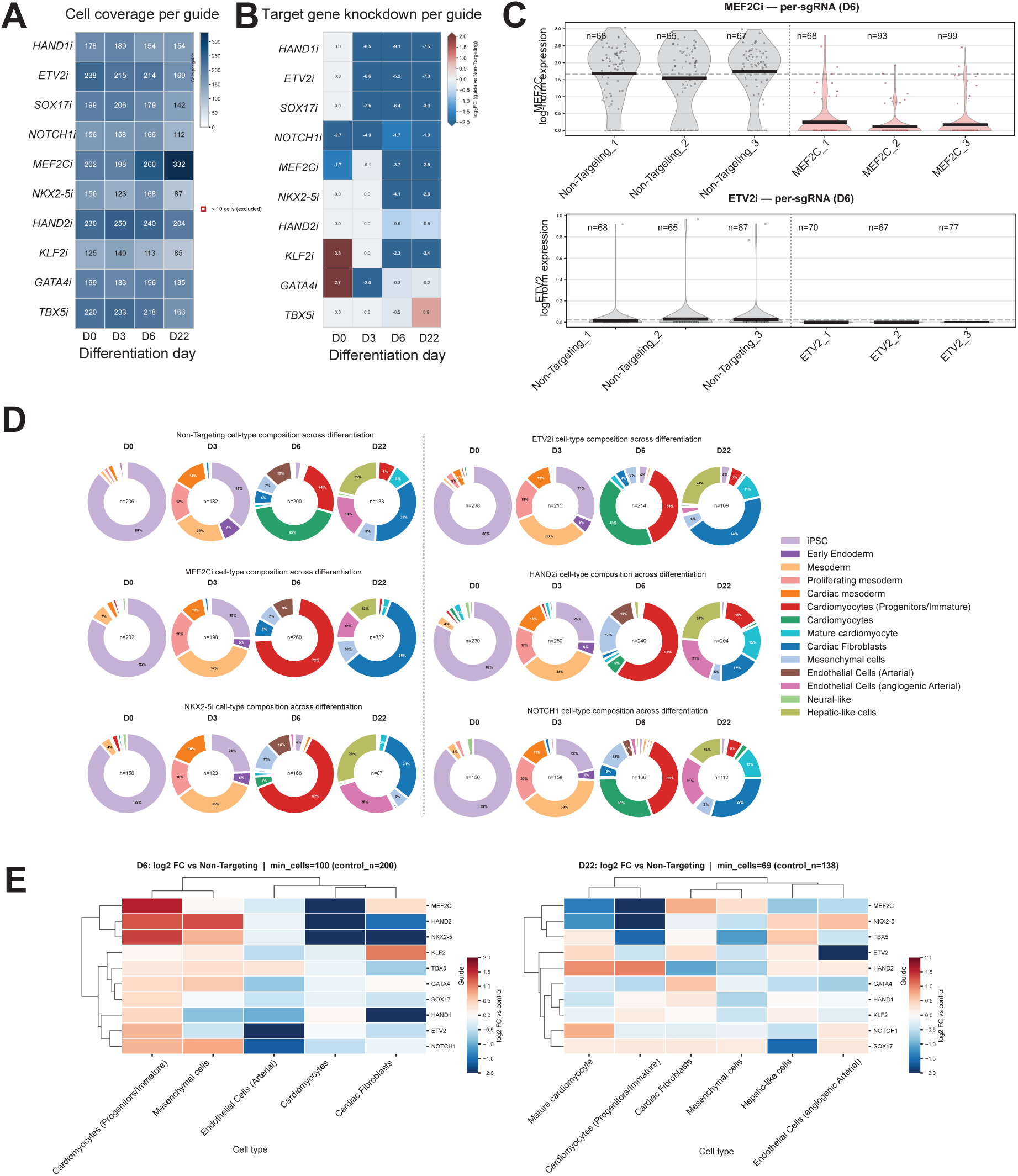
Stage- and lineage-specific effects of cardiovascular transcription factor perturbations. (A) Distribution of cells assigned to individual TF perturbations. (B) CRISPRi-mediated repression efficiency of target TFs across developmental stages. (D) Changes in cell-type proportions following TF knockdown at D6. (E) Hierarchical clustering of TF perturbations on D6 and D22 based on their effects on cell-type composition, highlighting distinct cardiac and vascular regulatory groups.

### A stage- and lineage-resolved phenotypic landscape for ten cardiovascular transcription factors

We quantified repression as the log2 fold-change of the target transcript in guide-assigned versus NTC cells, per day (**Fig. 2B**). Repression was strong and unambiguous for the two most highly expressed cardiac factors: MEF2C fell from a non-targeting mean of ≈1.37 to ≈0.19 log-normalised units at D6 (log2FC −3.3; Mann–Whitney p = 6.9 × 10⁻⁵³; n = 200 NTC vs 260 perturbed cells), and NKX2-5 from ≈1.12 to ≈0.09 (log2FC −3.8; p = 5.3 × 10⁻³⁵; n = 200 vs 168). All three sgRNAs behaved consistently for both genes. HAND2 transcript fell from ≈0.58 to ≈0.38 at D6 (log2FC −0.6; p = 4.2 × 10⁻³), and one of its three sgRNAs (HAND2_2) gave a higher mean than NTC. TBX5 was not significantly repressed at D6 (≈0.70 vs ≈0.65; p = 0.36), and GATA4 was reduced by roughly 7% (≈1.39 vs ≈1.29; p = 0.014; log2FC −0.2). For the four vascular factors and for HAND1, baseline expression was low but perturbation was significant (HAND1 p = 8.3 × 10⁻⁴; SOX17 p = 7.0 × 10⁻³; KLF2 p = 9.8 × 10⁻⁴; NOTCH1 p = 2.9 × 10⁻²; ETV2 p = 2.0 × 10⁻²).

Because the TFs were significantly activated in D6 around CPC stage and mostly highly perturbed around D6. We focused our analysis on D6 CPC stage, which also has diverse cell-type composition showing both cardiac and vascular lineage specification. Relative to NTC, MEF2C NKX2-5, and HAND2 silencing collapsed the cardiomyocyte compartment (log2FC ≈ −2 for both) with a reciprocal accumulation of cardiomyocyte progenitors/immature cells (log2FC ≈ +2), indicating an potential arrested transition rather than cell loss (**Fig. 2D**). NKX2-5, HAND1 and TBX5 silencing depleted cardiac fibroblasts (log2FC −1.3 to −2). ETV2 silencing almost completely depleted EC (log2FC ≈ −2), following with strong EC depletion effects by NOTCH1 silencing (≈ −1.5), while KLF2 and SOX17 produced modest effects at this stage. Hierarchical clustering of the compositional effects placed MEF2C, NKX2-5, and HAND2 together as a distinct CM perturbing group, while ETV2 and NOTCH1 as a EC perturbing group, separate from other TFs (**Fig. 2E**). D22 the compositional picture was consistent in direction. ETV2 silencing depleting angiogenic arterial endothelium (log2FC −2.6; 5 perturbed vs 25 control cells), SOX17 silencing depleting hepatic-like cells (−1.6; 10 vs 29), MEF2C silencing depleting mature cardiomyocytes (−1.4; 10 vs 11) and cardiomyocyte progenitors (−2.0; 6 vs 10), and HAND2 silencing depleting cardiac fibroblasts (−1.1; 34 vs 48) while enriching mature cardiomyocytes (+0.9; 31 vs 11) and progenitors (+1.0; 30 vs 10).

Because mRNA-level repression is a poor proxy for perturbation strength when the target is lowly expressed, we additionally computed a per-cell perturbation score in the manner of mixscape, which classifies cells by the magnitude of their perturbation signature rather than by target transcript abundance (**Fig. 3A**). Milo neighborhood-level differential abundance analysis revealed distinct lineage-specific effects of TF perturbations during organoid differentiation (**Fig. 3B**). Knockdown of MEF2C, HAND2, and NKX2-5 led to the accumulation of CPCs and depletion of differentiated cardiomyocytes, suggesting a developmental bottleneck at the CPC-to-cardiomyocyte transition. In contrast, ETV2 knockdown markedly depleted EC populations across developmental stages (D0–D22), highlighting its essential role in endothelial lineage specification.

**Figure 3.**
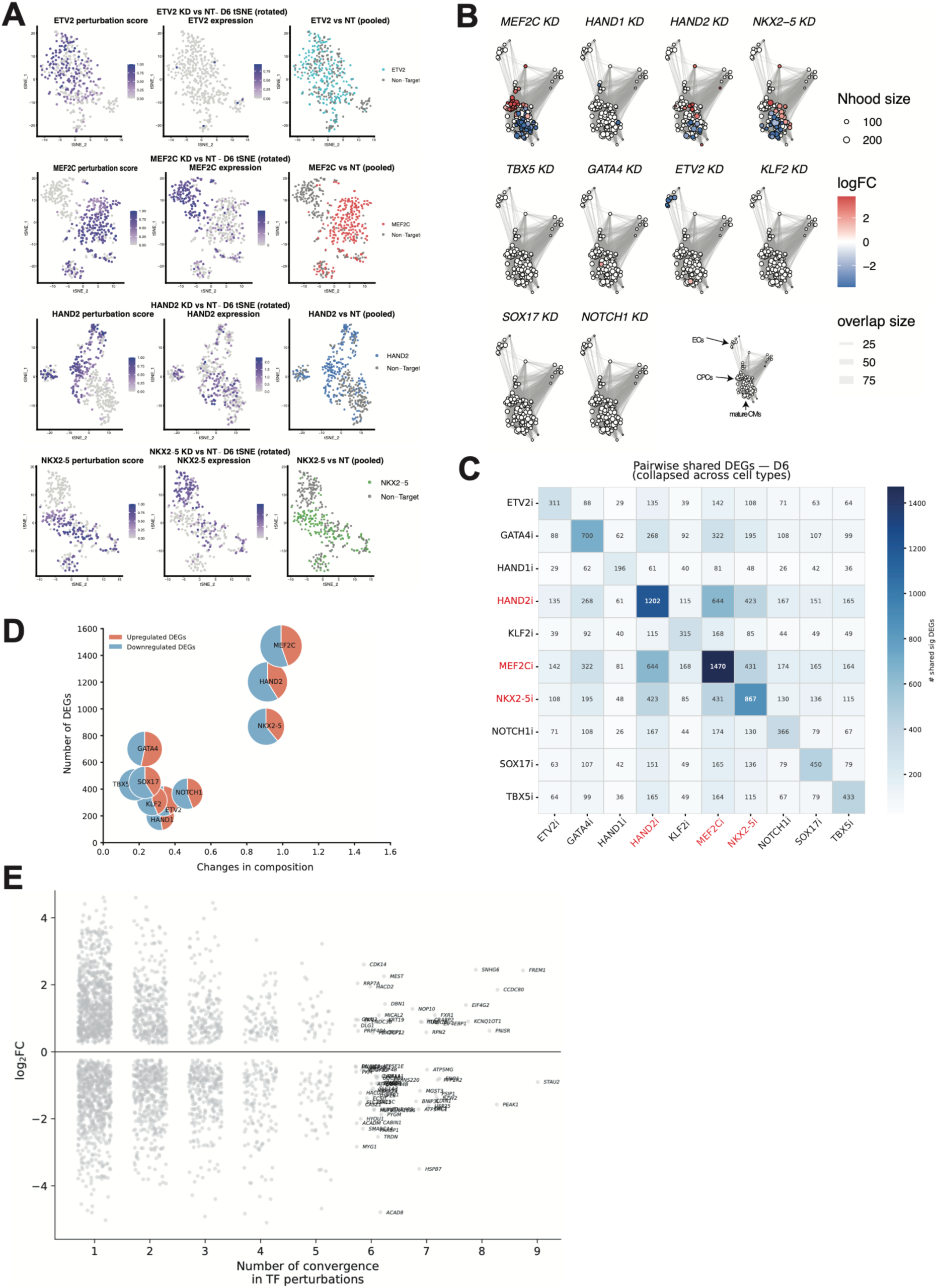
Perturbation-induced changes in cell fate and transcriptional programs. (A) Single-cell perturbation scores quantifying transcriptional responses to TF knockdown. (B) Milo neighborhood-level differential abundance analysis showing cardiac progenitor accumulation following MEF2C, HAND2, and NKX2-5 knockdown and endothelial depletion following ETV2 knockdown. (C–D) Comparison of cell-type compositional changes and differentially expressed genes (DEGs) across TF perturbations. (E) Shared transcriptional responses across multiple TF perturbations.

Plotting each perturbation by the magnitude of its cell type compositional shift against the number of differentially expressed genes (DEGs) it produced separated the library into two groups (**Fig. 3C-D**). MEF2C (1,344 DEGs), HAND2 (1,024) and NKX2-5 (754) formed a high-impact cluster with compositional shifts of ≈0.8–1.0, while GATA4 (645), SOX17 (449), TBX5 (398), NOTCH1 (340), ETV2 (297), KLF2 (292) and HAND1 (169) clustered at shifts below 0.5. Up- and down-regulated genes were close to balanced within each perturbation, arguing against a generic transcriptional shutdown. A small set responded to seven or more of the nine TF perturbations: FREM1 (up, log2FC +2.2, responding in 9), then STUB1, STAU2, PEAK1 and HSPB7 (all down, responding in 8), and SNHG6, MEST, CCDC80, HACD2, FXR1, CYP51A1, MLF1 and ISOC1 at 7 (**Fig. 3E**). Most of this high-convergence set is not cardiac-specific, and we hypothesize it as a shared cell-state response that accompanies cell fate perturbation of the D6 program rather than as a common direct target set.

### Convergent and divergent gene modules across transcription factor perturbations

To separate the shared from the TF-specific response we clustered perturbation log2FC profiles by k-means within each lineage. In D6 cardiomyocytes, 1,700 DEGs across the six cardiac TFs resolved into seven modules (**Fig. 4A**). Module 2 was coordinately up-regulated by MEF2C, HAND2 and NKX2-5 silencing and comprised cell migration, cell adhesion, locomotion and Wnt-signalling genes, with neurogenesis and head-development terms also enriched (GO BP, FDR < 0.05; cell migration k = 16, cell adhesion k = 15, regulation of Wnt signalling k = 7; Fig. 4E). Module 5 behaved differently: it was up-regulated by MEF2C silencing but down-regulated by HAND2 silencing, and its top Hallmark enrichment was epithelial–mesenchymal transition (k = 9, FDR q = 3 × 10⁻⁸), followed by myogenesis (k = 7), hedgehog signalling, hypoxia and angiogenesis (**Fig. 4A and 4C**).

**Figure 4.**
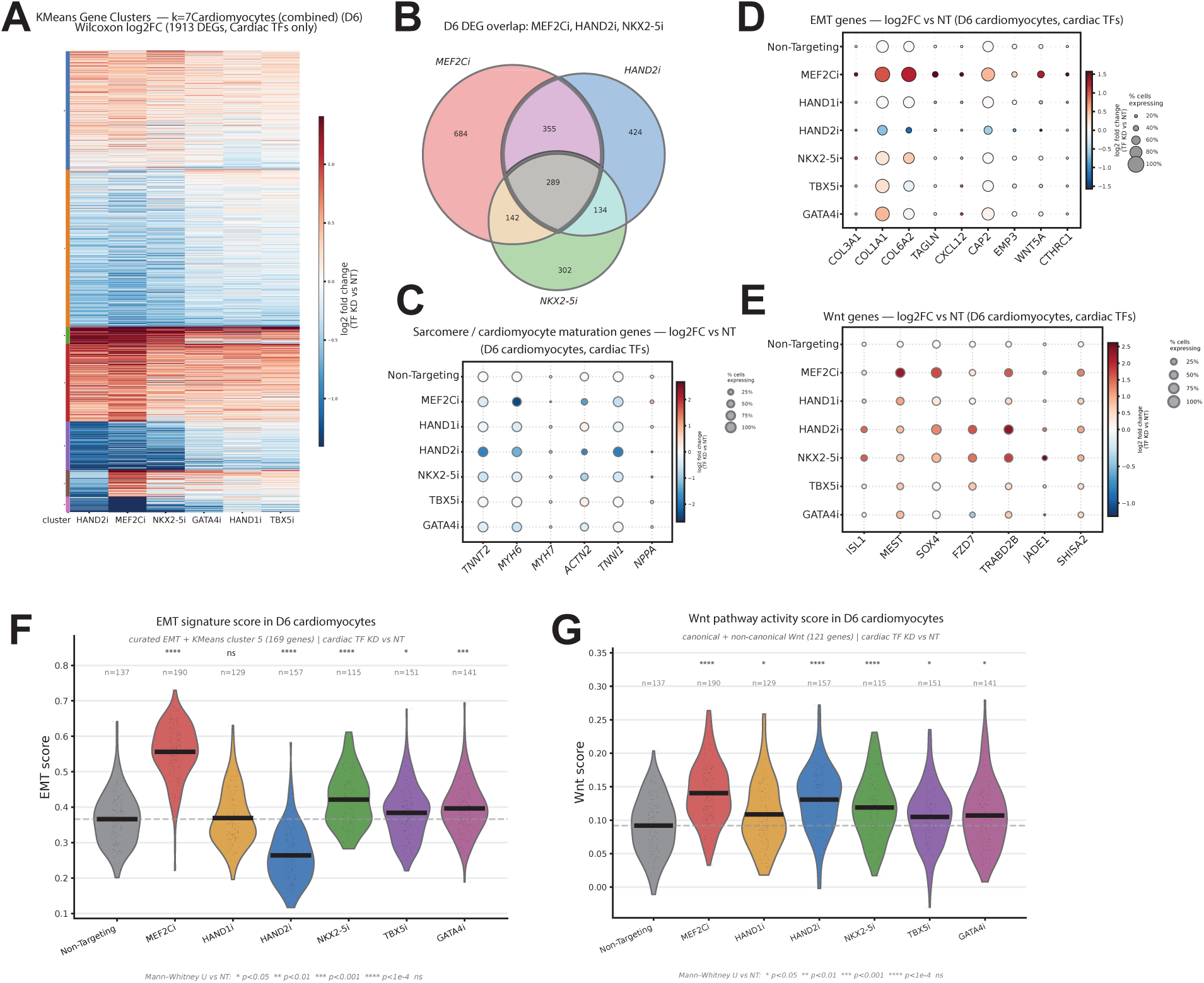
Convergent and divergent regulatory programs of MEF2C and HAND2 during cardiac differentiation. (A) Clustering of DEG modules following cardiac TF perturbations at D6. (B) Overlap of DEGs among MEF2C, HAND2, and NKX2-5 knockdown. (C) Functional enrichment of sacormere and maturaiton gene modules. (D) Comparison of MEF2C- and HAND2-dependent transcriptional responses, highlighting divergent regulation of mesenchymal genes. (E) Shared progenitor-associated and Wnt signaling programs. (F) EMT signature scores following cardiac TF perturbations. (G) Wnt pathway activity scores following cardiac TF perturbations

In D3 cardiac mesoderm, 451 DEGs across the four vascular TFs resolved into ten modules with a clear ETV2-specific block. ETV2 silencing alone produced 559 DEGs at D3 (278 down, 281 up), enriched for cellular stress response, cell-cycle and mitotic terms, RNA processing and splicing, and the E2F and MYC target Hallmark sets), a proliferative-arrest signature consistent with loss of the earliest endothelial specification factor before the endothelial program itself is established.

MEF2C and HAND2 are the two largest phenotypes in the screen, the most similar pair by shared DEGs, both second-heart-field-associated, and both required for cardiomyocyte production in mouse and zebrafish. We therefore examined the pair directly, with NKX2-5 as an internal comparator. A three-way intersection of D6 DEG sets defined a 491-gene MEF2C∩HAND2 core, of which 192 genes were also NKX2-5-responsive; 704 genes were MEF2C-specific, 419 HAND2-specific and 299 NKX2-5-specific (**Fig. 4B**).

### MEF2C and HAND2 converge on the progenitor-to-cardiomyocyte transition and divergently control the mesenchymal/EMT program

Convergence at the compositional level does not require convergence at every gene. Plotting MEF2C KD against HAND2 KD log2FC for all 1,318 D6 cardiomyocyte DEGs gave a positive but incomplete correlation (Pearson r = 0.56, Spearman ρ = 0.63; **Fig. 4D**). Most genes (n = 1,215) were convergent, but 81 genes were up-regulated by MEF2C silencing and down-regulated by HAND2 silencing, including NPPB, POPDC2, QRFPR, IGFBP7, S1PR1, BAALC and PDE3A, and 22 genes showed the opposite pattern, among them OPCML, GRID2, AFF3, GRIP1, PCLO, ZNF385B and RETREG1. Across a panel of EMT and matrix genes in D6 cardiomyocytes, COL3A1, COL1A1, COL6A2, TAGLN, CXCL12, CAP2, EMP3, WNT5A and CTHRC, MEF2C silencing raised mean expression and expressing fraction for every gene, while HAND2 silencing lowered both (**Fig. 4D**). NKX2-5, TBX5, GATA4 and HAND1 silencing produced intermediate or NTC-like profiles. Scoring each cell against a 159-gene EMT signature (k-means cluster 5 plus curated EMT genes) quantified the split (**Fig. 4F**). Median EMT score in NTC D6 cardiomyocytes was 0.295 (n = 137). MEF2C silencing raised it to 0.445 (n = 190; Mann–Whitney p < 10⁻⁴) and HAND2 silencing lowered it to 0.190 (n = 157; p < 10⁻⁴). NKX2-5 silencing also raised the score (0.353, n = 115; p < 10⁻⁴) and GATA4 silencing raised it marginally (0.322, n = 141; p < 0.05), while HAND1 (0.300, n = 129) and TBX5 (0.310, n = 151) were indistinguishable from control. MEF2C and HAND2 therefore sit on opposite sides of the control distribution for the same gene program in the same cell type at the same stage, despite producing the same compositional phenotype. We next asked whether this divergent transcriptional program translates into altered mesenchymal output at the tissue level. At D22, HAND2 silencing depleted the cardiac fibroblast compartment (log2FC −1.1; 34 perturbed vs 48 control cells) while MEF2C silencing did not (log2FC +0.7; 193 vs 48), the direction predicted by the D6 EMT scores.

The shared up-regulated program was progenitor- and Wnt-associated, indicating potential continuous activation of Wnt pathway impairing cardiac differentiation. ISL1, MEST, SOX4, FZD7, TRABD2B, JADE1 and SHISA2 were all expressed at higher mean level and in a larger fraction of D6 cardiomyocytes after MEF2C, HAND2 or NKX2-5 silencing than after NTC (**Fig. 4E**). ISL1 and FZD7 were most strongly elevated by HAND2 and NKX2-5 silencing; MEST and SOX4 by MEF2C silencing. The shared down-regulated program was the contractile apparatus: ACTN2 fell most steeply (log2FC ≈ −1.5 for MEF2C, ≈ −2.3 for HAND2), with MYH6 reduced by MEF2C, HAND2 and GATA4 silencing, and TNNT2 and TNNI1 reduced by HAND2 silencing (**Fig. 4C**). Taken together, cardiomyocytes lost, ISL1+ progenitors retained, Wnt and migration genes up, sarcomeric genes down, the data describe a failure to exit the multipotent progenitor state rather than a failure to survive as a cardiomyocyte (**Fig. 4G**). We summarize this as a convergent requirement: mesoderm gives rise to an ISL1+/APLNR+ multipotent progenitor, and MEF2C, HAND2 and NKX2-5 are each required for that progenitor to lock in the cardiac program, undergo metabolic maturation and express the contractile cardiomyocyte gene sets.

## Discussion

In this study, we established a time-resolved, multicellular Perturb-seq platform using human iPSC-derived cardiac organoids to systematically investigate TF functions during cardiovascular differentiation. By integrating CRISPRi perturbations with single-cell transcriptomic profiling across four developmental stages, we identified distinct stage- and lineage-specific requirements for key cardiac and vascular TFs. Perturbation of MEF2C, HAND2, and NKX2-5 resulted in the accumulation of cardiac progenitor cells and depletion of differentiated cardiomyocytes, suggesting a developmental bottleneck during cardiomyocyte commitment. In contrast, ETV2 and NOTCH1 perturbations preferentially impaired endothelial lineage specification, consistent with their established roles in vascular development. These findings demonstrate the utility of multicellular Perturb-seq for resolving transcriptional regulatory programs governing coordinated cardiac and vascular differentiation within a human organoid model.

Interestingly, although MEF2C and HAND2 perturbations produced similar defects in cardiomyocyte differentiation, they exhibited divergent effects on mesenchymal and epithelial–mesenchymal transition (EMT)-associated transcriptional programs. MEF2C silencing enhanced EMT-associated gene expression, whereas HAND2 silencing suppressed this program, suggesting that these TFs regulate distinct aspects of cardiac lineage specification beyond their shared roles in cardiomyocyte commitment. These findings highlight the complexity of TF regulatory networks, in which individual factors can exert convergent effects on cell fate while differentially regulating alternative lineage programs. Future studies integrating combinatorial perturbations, lineage tracing, and functional validation will be important to establish the underlying regulatory mechanisms and determine how cardiac and vascular TFs coordinate multicellular interactions during cardiogenesis. Furthermore, extending this platform to disease-associated genes and genetic variants could provide a systematic approach to identifying disrupted regulatory networks underlying congenital cardiovascular disorders.

## Methods

### Human iPSC differentiation and generation of cardiac organoids

Human iPSCs harboring a dCas9-KRAB CRISPR interference (CRISPRi) system were differentiated into cardiac organoids (COs) using an optimized protocol adapted from previously published cardiac organoid and gastruloid differentiation methods. Briefly, mesoderm differentiation was initiated at day 0 (D0), followed by cardiac patterning at D3-D4. Vascular differentiation was promoted by the addition of vascular endothelial growth factor (VEGF) at D4. The resulting COs were maintained until D22, during which cardiomyocyte and endothelial populations differentiated and self-organized within the same organoids. Cells were collected at D0, D3, D6, and D22 for single-cell transcriptomic analysis to investigate transcription factor functions across different stages of cardiovascular differentiation.

### CRISPRi sgRNA library and pooled genetic perturbation

A pooled CRISPRi sgRNA library was designed to target six cardiac transcription factors (GATA4, HAND1, HAND2, MEF2C, NKX2-5, and TBX5) and four vascular transcription factors (ETV2, KLF2, NOTCH1, and SOX17). Three sgRNAs were included for each target gene, together with three non-targeting control (NTC) sgRNAs. Human CRISPRi iPSCs were transduced with the pooled lentiviral sgRNA library at a multiplicity of infection (MOI) of 0.2–0.3 to minimize multiple sgRNA integrations per cell. To avoid potential effects of antibiotic selection on iPSC maintenance and subsequent differentiation, transduced iPSCs were not subjected to puromycin selection. The transduced iPSCs were differentiated into COs, and cells were harvested at D0, D3, D6, and D22 for simultaneous identification of sgRNAs and single-cell transcriptomic profiling.

### Single-cell data sequencing and data preprocessing

Single-cell gene expression and CRISPR guide capture data were generated using the 10x Genomics Single Cell 5′ platform. Raw sequencing data from each differentiation time point (D0, D3, D6 and D22) were processed with Cell Ranger (v9.0.1; 10x Genomics) against the GRCh38-2024-A human reference (Single Cell 5′ PE v3 chemistry), producing gene expression and CRISPR guide capture count matrices. Low-quality cells were identified with a median absolute deviation (MAD) based outlier approach. Cells were flagged if they deviated by more than 5 MADs from the median in log-transformed total UMI counts, log-transformed number of detected genes, or the percentage of counts in the top 20 most highly expressed genes. Cells were additionally flagged if their mitochondrial read fraction deviated by more than 3 MADs or exceeded 8%. Doublets were identified with scDblFinder (v1.24.0). Cells flagged as outliers, as high-mitochondrial, or classified as doublets were removed. This retained 8,613 of 13,663 cells at D0, 7,831 of 13,807 at D3, 8,546 of 12,788 at D6 and 5,168 of 6,870 at D22 (30,158 cells in total). Ambient RNA contamination was corrected with SoupX (v1.6.2). The ambient soup profile was estimated from droplets in the raw, unfiltered matrix containing fewer than 10 UMIs. Because automated contamination estimation performed poorly on these data, the contamination fraction was fixed at 0.05, and corrected counts were obtained with adjustCounts. SoupX-corrected counts were used as the raw count input for all downstream analyses.

### sgRNA assignment and perturbation identification

sgRNA identities were assigned from the CRISPR guide capture library using the per-protospacer UMI thresholds computed by Cell Ranger. A cell was assigned a guide if its UMI count for that guide met or exceeded the guide-specific threshold. The number of guides passing threshold was recorded for each cell. Only cells carrying exactly one assigned guide were retained for downstream analysis, yielding 8,068 cells across all four time points. Individual guides were collapsed to their target gene (for example, HAND1_1, HAND1_2 and HAND1_3 to HAND1), and cells carrying non-targeting guides served as controls.

### Single-cell data integration, clustering, and cell type annotation

Single-guide cells from the four time points were concatenated, and CRISPR guide features were removed. The top 3,000 highly variable genes were selected from the SoupX-corrected counts with the Seurat v3 method, using differentiation day as the batch key. A single-cell variational inference (scVI) model was trained on these genes with scvi-tools (v1.1.6.post2), with raw counts as input, differentiation day as a categorical covariate, and a 30-dimensional latent space. Default architecture and training settings were used, and the model was trained for 400 epochs. All random seeds (Python, NumPy, PyTorch and scvi-tools) were set to 0, and deterministic computation was enforced for reproducibility.

A 15-nearest-neighbor graph was constructed on the scVI latent space and used for Leiden clustering (resolution 2.0) and UMAP embedding. The resulting 27 clusters were annotated into 14 major cell states based on canonical lineage markers, including POU5F1, SOX2, and NANOG for pluripotent cells; TBXT, EOMES, MIXL1, MESP1, KDR, and PDGFRA for mesoderm populations; ISL1, NKX2-5, GATA4, TBX5, MEF2C, and HAND2 for cardiac progenitors; TNNT2, ACTN2, MYH6, MYH7, TNNI3, and RYR2 for cardiomyocytes; PECAM1, CDH5, VWF, and CLDN5 for endothelial cells; COL1A2, DCN, LUM, and POSTN for fibroblasts; SOX10, TFAP2A, and FOXD3 for neural crest-associated populations; and ALB, SERPINA1, and HNF4A for hepatic-like populations. Random seeds were fixed to ensure reproducibility of data integration and clustering.

### Cell type composition and differential abundance analysis

Cell-type composition was summarized for each target and day as the fraction of cells assigned to each cell type. Cell types that never exceeded 0.5% of cells at any day were omitted from composition plots. Compositional shifts were quantified as the log2 ratio of the fraction of perturbed cells in a cell type to the corresponding fraction of NTC cells from the same day, log2((f_KD + 0.0001) / (f_NTC + 0.0001)). Fractions were calculated over all cells of a given target and day and then reported for the same predefined major cell types used for knockdown estimation. Targets were included at a given day only if they had at least max (20, 0.5 × the number of NTC cells at that day) cells.

Perturbation-associated shifts in cell state at D6 and D22 were assessed with Milo (miloR v2.6.0), which tests for differential abundance in overlapping neighborhoods of a k-nearest-neighbor graph. The processed AnnData object was imported into R with zellkonverter (v1.20.1), and cells from each day were analyzed separately. A k-nearest-neighbor graph (k = 30) was built on the 30-dimensional scVI latent space, and refined neighborhoods were sampled from 10% of cells. Cells were counted per neighborhood with each sgRNA treated as a sample, so that the three guides against each target acted as pseudo-replicates. For each target, neighborhood counts from the target’s sgRNAs were compared with those from the NTC sgRNAs using testNhoods with the design ∼ condition, and targets with fewer than two sgRNAs on either side were skipped. Because sgRNAs are technical rather than biological replicates, spatial FDR values were not treated as formal inferential evidence. Results were interpreted through the magnitude and direction of neighborhood log fold changes, which were displayed on the UMAP-based neighborhood graph with a symmetric color scale capped at the 95th percentile of absolute log fold change across all targets (minimum 1). Neighborhoods were shaded at a spatial FDR threshold of 0.1 at D6 and 0.5 at D22.

### Perturbation scores

Per-cell perturbation scores at D6 were estimated with scMAGeCK (v1.9.2). Perturbed cells and NTC cells were extracted from a Seurat (v5.5.0) object built from the D6 counts, which were log-normalized, subset to variable features, and scaled with default settings. Perturbation scores were computed with scmageck_eff_estimate and visualized on a t-SNE embedding computed from the first 30 principal components, alongside target-gene expression. Scores were summarized per sgRNA (mean, median and standard deviation) to assess consistency between guides. The random seed was set to 1.

### Differential expression, gene module identification, and pathway enrichment analysis

Differential expression was performed on D6 cells, using log1p CP10k values recomputed from SoupX-corrected counts. To retain power, the three cardiomyocyte populations (progenitor/immature, cardiomyocytes and mature cardiomyocytes) were pooled into a single cardiomyocyte group. Within each cell type, cells carrying guides against each target were compared with NTC cells using the two-sided Wilcoxon rank-sum test implemented in Scanpy rank_genes_groups. Genes were called differentially expressed at nominal P < 0.05 and 0.25 < |log2 fold change| < 10, with the upper bound excluding unstable estimates from near-zero expression.

To identify gene programs co-regulated by the cardiac transcription factors, the union of DEGs from the six cardiac targets (MEF2C, HAND1, HAND2, NKX2-5, TBX5 and GATA4) in pooled D6 cardiomyocytes was assembled. A gene × target matrix of Wilcoxon log2 fold changes was built from the full test results. Genes were clustered with k-means (k = 10, 20 initializations, random state 42), and clusters with fewer than 20 genes were merged into the cluster with the nearest centroid by Euclidean distance. Targets were ordered by hierarchical clustering with average linkage on correlation distance. We run GO enrichment on each cluster gene and idenfity the top terms associated with each cluster of genes by using GSEA. Pathway-level activity was then scored in individual D6 cardiomyocytes with Scanpy score_genes (random state 42). Gene sets comprised an epithelial–mesenchymal transition (EMT) signature, combining curated EMT genes drawn from the MSigDB Hallmark EMT set and the literature with the genes of k-means cluster 5, and a combined canonical and non-canonical Wnt signaling set. Scores for each cardiac target were compared with NTC cells using two-sided Mann–Whitney U tests.

### Software and statistical analysis

Analyses in Python (v3.9.6) used Scanpy (v1.10.3), AnnData (v0.10.8), scvi-tools (v1.1.6.post2), PyTorch (v2.8.0), leidenalg (v0.11.0), scikit-learn (v1.6.1), SciPy (v1.13.1), pandas (v2.3.3), NumPy (v1.26.4), seaborn (v0.13.2) and Matplotlib (v3.9.4); R (v4.5.2) analyses used scDblFinder (v1.24.0, xgboost v3.1.2.1), SoupX (v1.6.2), miloR (v2.6.0), scMAGeCK (v1.9.2), Seurat (v5.5.0), SingleCellExperiment (v1.32.0), zellkonverter (v1.20.1) and ggplot2 (v4.0.3), called from Python through rpy2 (v3.6.7) and anndata2ri (v1.3.2) where noted. Random seeds were fixed for every stochastic step as stated above.

### Availability of data and materials

Data will be uploaded to GEO dataset.

